# Antifungal effects of alantolactone on *Candida albicans:* an *in vitro* study

**DOI:** 10.1101/2021.01.25.428193

**Authors:** Xin Liu, Lili Zhong, Zhiming Ma, Yujie Sui, Jia’nan Xie, Tonghui Ma, Longfei Yang

## Abstract

The human fungal pathogen *Candida albicans* can cause many kinds of infections, including biofilm infections on medical devices, while the available antifungal drugs are limited to only a few. In this study, alantolactone (Ala) demonstrated antifungal activities against *C. albicans*, as well as other *Candida* species, with a MIC of 72 μg/mL. Ala could also inhibit the adhesion, yeast-to-hyphal transition, biofilm formation and development of *C. albicans*. The exopolysaccharide of biofilm matrix and extracellular phospholipase production could also be reduced by Ala treatment. Ala could increase permeability of *C. albicans* cell membrane and ROS contribute to the antifungal activity of Ala. Overall, the present study suggests that Ala may provide a promising candidate for developing antifungal drugs against *C. albicans* infections.

## Introduction

An important threat to human health is fungal infections, among which infections caused by *C. albicans* are one of the most common (Brown et al., 2012; Perlin et al., 2017). Although opportunistic, this fungus could infect most organs of mammals with compromised immunity, causing oropharyngeal and esophageal candidiasis, and vaginitis (Mayer et al., 2013). In the clinical context, *C. albicans* can cause life-endangering candidemia and dwell on medical devices, such as catheters, leading to the formation of drug-resistant biofilms, where the exopolysaccharide (EPS) of extracellular matrix, the increased cell density, overexpression of drug targets, upregulation of drug efflux pumps and the presence of persister cells, contributes to the antifungal resistance (Fox and Nobile, 2014; Liu et al., 2017; Wall et al., 2019). The calcitrant *C. albicans* biofilms, along with the drug resistance and the undesired side effects of the handful of currently available antifungal drugs, call for the development of new antifungal agents (Liu et al., 2017; Perlin et al., 2017).

Natural products, especially those from traditional medicines, are promising candidates for antifungal therapies (Liu et al., 2017). Alantolactone (Ala), the major active sesquiterpene lactone component of *Inula helenium*, is a pleiotropic molecule showing anti-proliferative activities against various kinds of tumors, such as hepatocarcinoma (Khan et al., 2013), leukemia (Yang et al., 2013), lung adenocarcinoma (Zong et al., 2011; Maryam et al., 2017), and glioblastoma (Khan et al., 2012). Other pharmacological activities include neuroprotective (Wang et al., 2018), antiviral (Rezeng et al., 2015), and anti-inflammatory (Kim et al., 2017), as well as antibacterial activities against *Mycobacterium tuberculosis* and *Staphylococcus aureus* (Cantrell et al., 1999; Stojanovic-Radic et al., 2012).

In addition, Ala has also demonstrated fungistatic activity against *Fusarium solani, in vitro*, a fungal pathogen for plants and human, at concentrations above 100 μg/mL (Wahab et al., 1979). However, the antifungal activities of Ala against *C. albicans*, has never been explored. Therefore, this study was conducted to explore the activity of Ala against *C. albicans*, as well as the underlying mechanisms.

## Materials and methods

In this study, the widely-used strain *C. albicans* SC5314 was selected for its capacity to form biofilms. Several other *Candida* species, namely *C. albicans* ATCC10231, *Candida glabrata* ATCC2001, *Candida krusei* ATCC6258, and *Candida tropicalis* ATCC7349 were also used to evaluate the antifungal activity of Ala. All these strains were bought from CGMCC and grown on YPD (yeast extract-peptone-dextrose) agar. Before each test, fungal cells were propagated in YPD medium at 28°C overnight with a rotation of 150 rpm.

Ala, Amphotericin B (AmB), fluconazole and caspofungin acetate (CAS) were bought from Solarbio, 2,3-bis(2-methoxy-4-nitro-5-sulfophenyl)-2H-tetrazolium-5-carboxanilide (XTT), Calcofluor White (CFW), N-Acetyl-glucosamine (GlcNAc), N-acetyl-cysteine (NAC), propidium iodide (PI) and 2′,7′-Dichlorodihydrofluorescein diacetate (DCFH-DA) were bought from Sigma Aldrich (Shanghai). Ala and antifungal drugs were dissolved in DMSO to get stock solutions of 20 mM.

### Antifungal susceptibility tests

The antifungal susceptibility of Ala was assessed in RPMI-1640 medium through micro-dilutions following the guidelines of CLSI-M27-A3, with little modifications (Yang et al., 2018c). The concentration at which no fungal growth was observed was defined as minimum inhibitory concentration (MIC). After MIC detection, entire cultures in each well containing higher concentrations of Ala than MIC were plated on YPD agars and incubated for 2 days. The lowest concentration at which no fungal colony grown on YPD agars was defined as minimum fungicidal concentration (MFC).

### Adhesion assay

This assay was performed to assay the influence of Ala on the adhesion of *C. albicans* to polystyrene surfaces of 96-well plates (Li et al., 2015). 100 μL of *C. albicans* suspension (10^6^ cells/mL in 1640 medium) was added in each well and after treatment with different concentrations of Ala for 90 minutes and PBS washing, the viability of fungal cells left in each well was determined by XTT assay to calculate the percentage of adherent cells.

### Growth rate determination

To assess the growth of *C. albicans* in the presence of Ala, overnight grown cultures were subcultured into YPD medium at a density of 10^6^ cells/mL, and incubated with 18, 36 and 72 μg/mL of Ala for 24 h at 37 °C, 140 rpm. 100 μL from such culture was transferred into 96-well plates for OD_600_ determination at intervals of two hours. These assays were performed in triplicate and repeated for three times.

### Yeast-to-hyphal transition

The influence of Ala on morphological transition of *C. albicans* was assessed in four hyphal-inducing media, RPMI-1640, Spider medium, GlcNAc medium and 10% FBS SD medium. 0, 18, 36 and 72 μg/mL of Ala were added into *C. albicans* cell suspensions (10^6^ cells/mL) in each medium and incubated for 4 hours at 37’C. Then, cellular morphologies were recorded by inverted microscope (IX71).

### Biofilm assay

*C. albicans* cell suspension (10^6^ cells/mL, in 1640 medium) was added into 96-well plates and allowed to grow statically to form biofilms in the presence of different concentrations of Ala for 24 h. After washing with PBS, the viability of cells in each well was determined by XTT assay (Siles et al., 2013). To assess the effects of Ala on biofilm development, mature biofilms formed without drugs were washed with PBS, and treated with Ala (in fresh 1640 medium) for another 24 h. Then, XTT reduction assay were performed.

### CLSM analysis

Biofilms, formed under exposure to different concentrations of Ala, were stained with CFW (50 μg/mL), and subjected to confocal laser scanning microscope (CLSM) to observe the structures of biofilms. Pictures acquired through xyz mode of scanning were reconstructed with Imaris 7.02 software.

### EPS detection

The EPS of preformed biofilms was determined by colorimetry as described by others (Nithyanand et al., 2015). Preformed *C. albicans* biofilms in 24-well plates were treated by different concentration of Ala for 24 h and washed with 0.9% NaCl solution. Then, 200 μL 0.9% NaCl, 200 μL 5% phenol and 2 mL 0.2% hydrazine sulfate (dissolved in sulfuric acid) were added into each well and plates were kept in dark for 1 h. The absorbance of reaction product in each well was detected at 490 nm by a multifunctional microplate reader (VarioSkan, Thermo).

### Extracellular phospholipase assay

The production of extracellular phospholipase was evaluated by egg yolk emulsion agar (Padmavathi et al., 2015). 1 μL of *C. albicans* cell suspension (10^6^ cells/mL) was added on agars supplemented with different concentrations of Ala, followed by incubation at 37’C for 4 days. The diameter of colony (*d*_*1*_), as well as that of colony and precipitation zone (*d*_*2*_) was measured. Pz (*d*_*1*_/*d*_*2*_) was used to assess the production of enzyme, while a bigger Pz stands for weak enzymatic production.

### PI influx

*C. albicans* cells (10^6^ cells/mL in 1640 medium) were incubated with different concentrations of Ala at 37°C for 4 h before cells were stained with 10 μM PI, a fluorescent dye that gains access into cells when cell membrane was damaged. After incubation in dark for 10 minutes, cells were subjected to flow cytometry (FCM) (Beckman Coulter, EPICS XL-MCL, US) to quantify the cells stained by PI.

### ROS detection

To determine the ROS production induced by Ala in *C. albicans* cells, DCFH-DA staining (10 μM, 30 minutes) were performed after fungal cells (10^6^ cells/mL in 1640 medium) were treated with different concentrations of Ala for 4 h at 28°C, 150 rpm. Then, fungal cells were washed with PBS and subjected to FCM for quantitative analysis.

### NAC rescue assay

To further test the effects of oxidative stress caused by Ala on the biofilm formation, N-acetyl-cysteine (NAC) rescue assay was performed. Under biofilm formation conditions, fungal cells in 1640 medium supplemented with or without 5 mM NAC was challenged with 32 μg/mL of Ala for 24 h. Then, the morphologies of biofilms were photographed before XTT assay were performed to assay the viability of cells in biofilms (Yang et al., 2018a).

### Checkerboard assay with antifungal drugs

To elucidate the potential interactions between Ala and antifungal drugs (AmB, fluconazole and CAS), checkerboard assays were performed (Haque et al., 2016). The interaction of combination was considered as synergistic when FICI is ≤ 5, additive when FICI is >0.5 and ≤ 1, indifferent when FICI is > 1 and ≤ 4, and antagonist when FICI is > 4.

### Statistical analysis

Data (expressed as mean + SD, from at least three independent assays) were analyzed with GraphPad Prism 6.02 (GraphPad Software, USA) and statistical significance was determined by Student *t*-test, with * indicating *P*<0.05.

## Results

From the results of CLSI microdilution assays, as shown in Table 1, the MIC of Ala against *C. albicans* SC5314 was 72 μg/mL, while the MFC of also 72 μg/mL, yielding an MFC/MIC ratio of 1, which indicated a fungicidal activity of Ala. The antifungal activities of Ala against other *Candida* species were similar to that against *C. albicans* SC5314. Therefore, the widely-used standard strain *C. albicans* SC5314 was selected for further assays in this study.

**Table 1.**
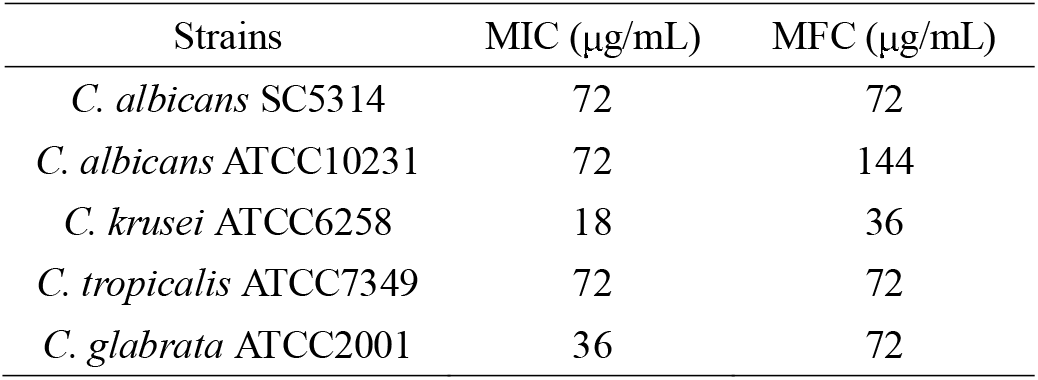
Antifungal susceptibility of Ala against *Candida* species.

Hyphal formation is closely associated with biofilm formation and escape from immune surveillance. As shown in Figure 1, with the increase of Ala concentration, the hyphal growth of *C. albicans* in all four kinds of media was inhibited gradually. The degree of inhibition was a little different, with inhibition in Spider medium be strongest where Ala at 36 μg/mL could keep cells in yeast type. Although hyphal induction in medium containing FBS could be suppressed by 72 μg/mL of Ala, the cells in yeast type were more than those in other media, indicating the strong induction of FBS and the weak inhibition of Ala in this FBS-containing medium.

**Figure 1.**
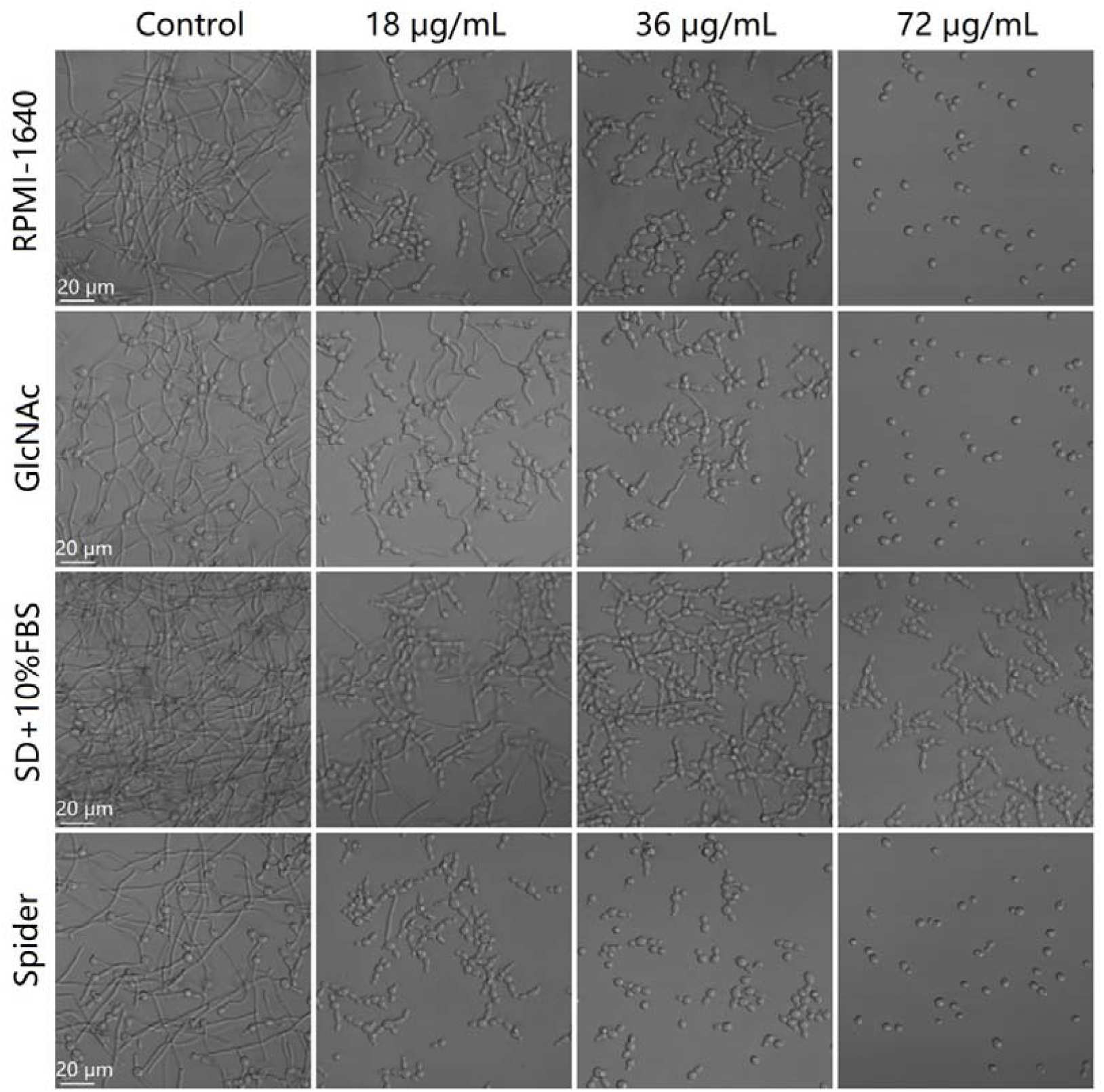
Hyphal induction in four kinds of media, namely RPMI-1640, GlcNAc, SD+10% FBS and Spider medium, was impeded by Ala in a concentration-dependent manner. After inoculation into each fresh medium to achieve a density of 10^6^ cells/mL, *C. albicans* cells were incubated at 37°C for 4 h, followed by microscope inspection and micrograph capture through inverted microscope (Olympus IX 71, Japan).

The widely used XTT reduction assay was performed to determine the antibiofilm activity of Ala. At concentrations below MIC, namely 9, 18, 36 and 72 μg/mL, Ala could significantly inhibit the biofilm formation of *C. albicans*, with a half maximal inhibitory concentration (IC_50_) of about 9 μg/mL (Figure 2A). 18, 36 and 72 μg/mL of Ala could also reduce the metabolic activity of preformed biofilm by 20%, 65% and 85% respectively, as compared to drug-free controls (Figure 2B). Adhesion plays important roles in biofilm formation and infections of *C. albicans*. The inhibitory effect of Ala on the adhesion of *C. albicans* cells to polystyrene surfaces was also quantified by XTT assay. As shown in Figure 2C, 18, 36 and 72 μg/mL of Ala could also significantly decrease the adhesion to polystyrene surfaces, to an extent of 15%, 40% and 88%, respectively.

**Figure 2.**
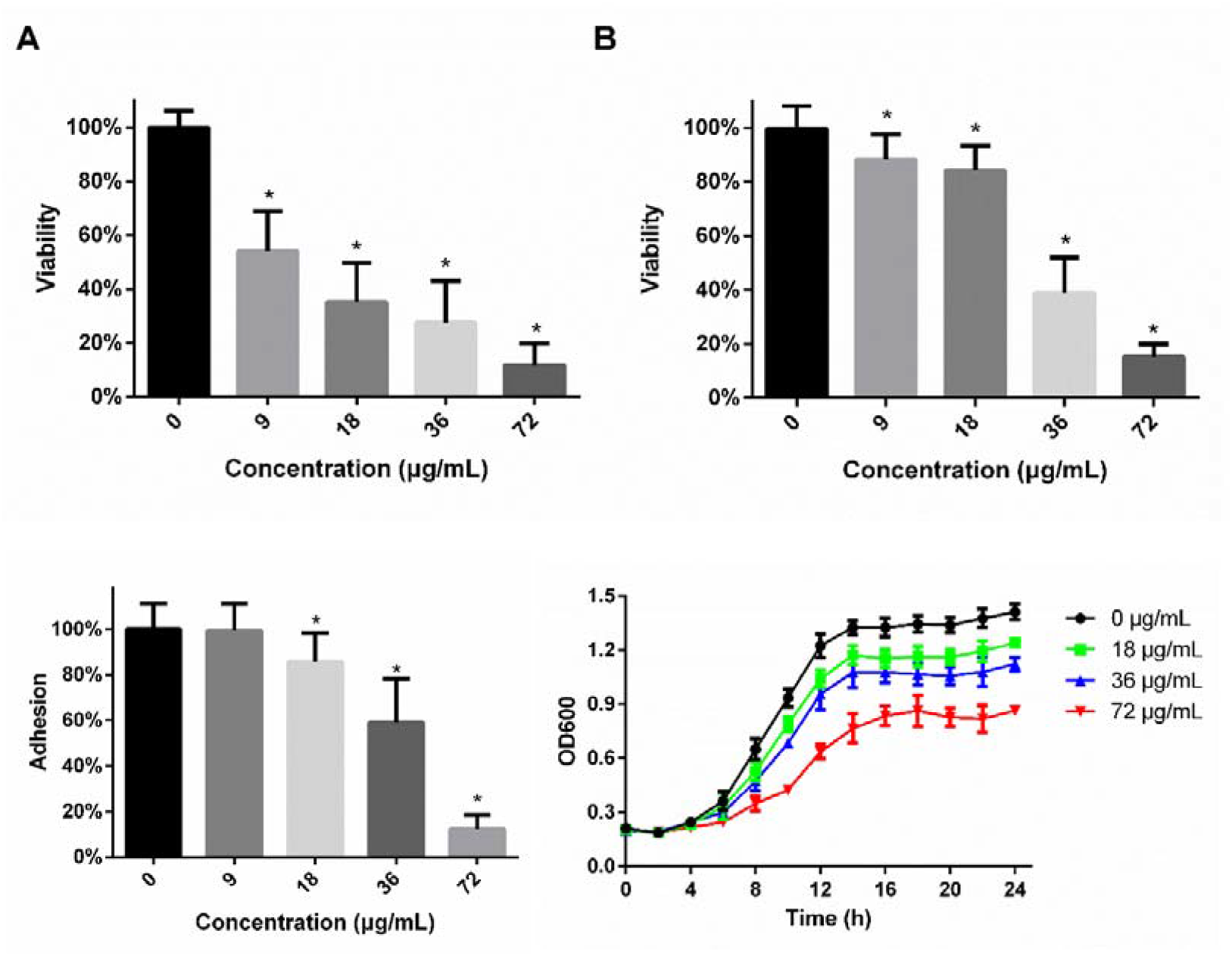
The effects of Ala on the biofilm formation and development. *C. albicans* biofilms were formed under exposure to Ala (A) or preformed biofilms were treated by Ala (B) for 24 h, followed by XTT reduction assay. (C) Ala decreased the adhesion of *C. albicans* cells to polystyrene surfaces. Adhesion to polystyrene surfaces was determined by XTT, which was used to quantify the viability of the residual cells standing 90 minutes of Ala treatment and subsequent PBS washing. *, p<0.05, Ala-treated vs control. (D) 18, 36 and 72 μg/mL of Ala did not inhibit completely the growth of *C. albicans*.

The inhibition of Ala on biofilm formation could also be confirmed by results from CLSM. Pictures of biofilms formed under treatment of Ala were recorded by CLSM and reconstructed by software to visualize the 3D structures. As shown in Figure 3, increasing the Ala concentration from 0 to 72 μg/mL, lead to the thinner and sparser biofilms. While treatment with 72 μg/mL of Ala can almost completely block the biofilm formation, and only yeast cells can be seen.

**Figure 3.**
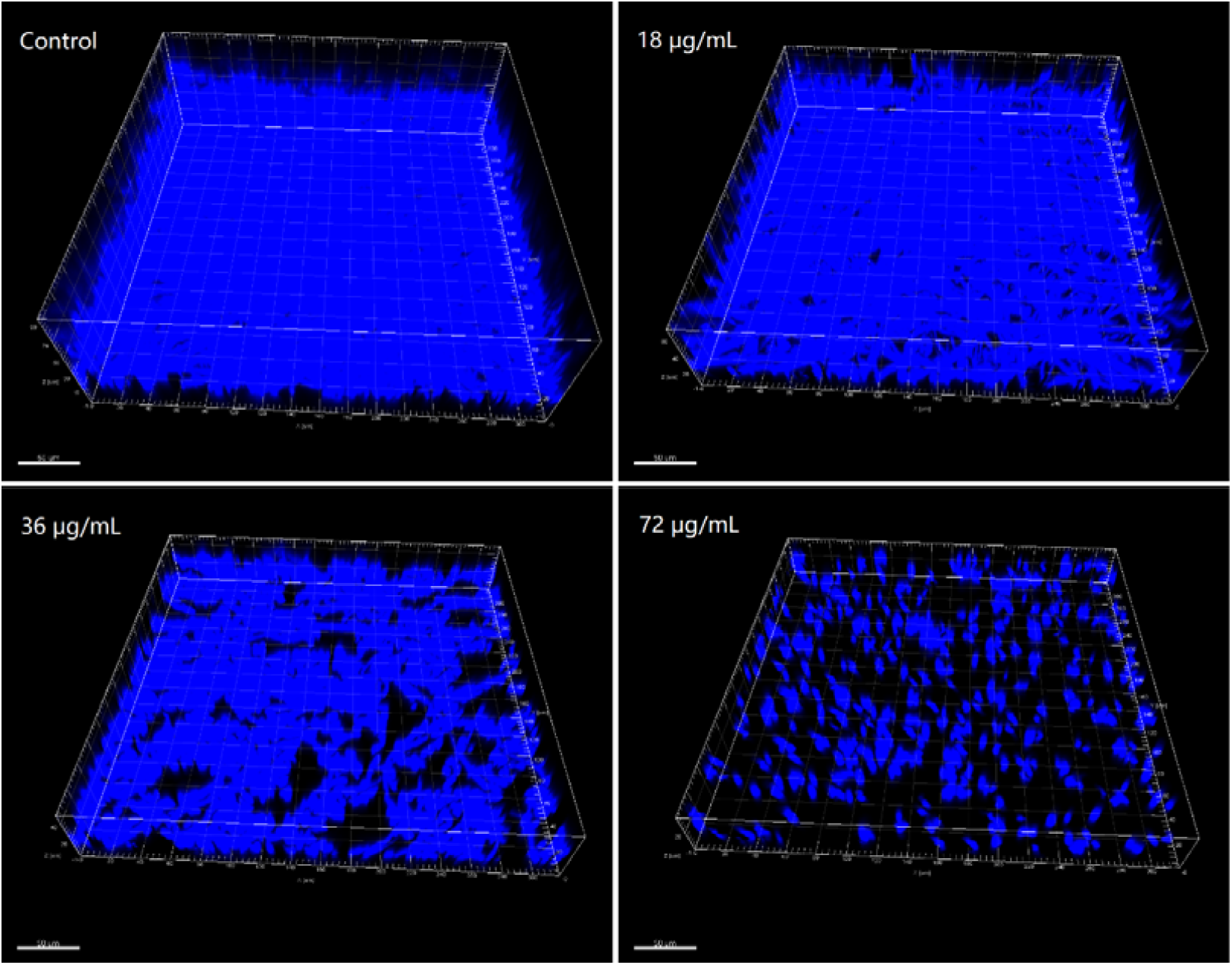
Ala inhibited biofilm formation, visualized and recorded by confocal microscope. Cells in biofilms were stained with 20 μg/mL CFW before microscope analysis. *Z*-stacked images obtained by CLSM were re-constructed by Imaris software version 7.02.

EPS of preformed *C. albicans* biofilm contribute to the sequestration of antifungal drugs within biofilms. So, we tested the effects of Ala on EPS production of preformed biofilms. As shown in Figure 4, Ala treatment could decrease the EPS production in mature biofilms, in a concentration-dependent way. Treatment with 18, 36 and 72 μg/mL of Ala could reduce the EPS production about 15%, 30% and 34%, as compared to drug-free controls.

**Figure 4.**
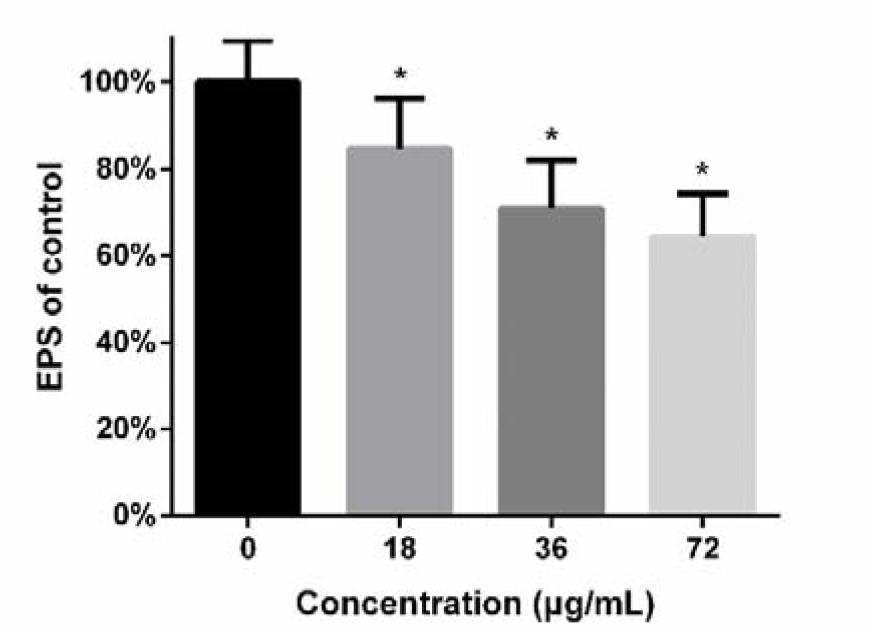
Ala inhibits the EPS production of *C. albicans* biofilms. Preformed *C. albicans* biofilms were treated with different concentrations of Ala for 24 h before the determination of EPS production in biofilms through phenol-hydrazine sulfate method. *, p<0.05 vs control.

Extracellular phospholipase of *C. albicans* contributes to its virulence in mouse models. Our results from egg yolk emulsion agar assay showed that Ala could significantly inhibit the production of extracellular phospholipase of *C. albicans* (Figure 5), although the increased extent of Pz was limited, which was considered as an indicator of phospholipase production.

**Figure 5.**
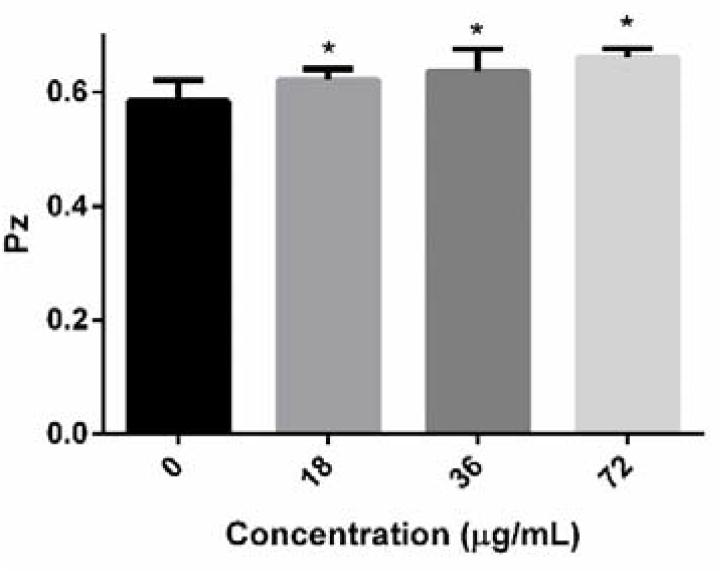
Ala affected the production of extracellular phospholipase in *C. albicans*. Fungal cells were added on egg yolk emulsion agars supplemented with different concentrations of Ala. After 4-day incubation at 37°C, the diameter of colony (*d*_*1*_) and that of colony plus precipitation zone (*d*_*2*_) were determined. Pz means *d*_*1*_/*d*_*2*_, while higher Pz indicates lower phospholipase production. *, p<0.05, Ala-treated vs control

After we found the inhibitory effects of Ala on the virulence factors of *C. albicans*, we proceeded to explore the mechanisms underlying the antifungal activity of Ala. At first, the effects of Ala on the permeability of *C. albicans* plasma membrane were determined by FCM after treated cells were stained by PI. Treatment with Ala for 4 hours, could increase the membrane permeability of *C. albicans* cells, as evidenced by increased proportion of cells stained by PI (Figure 6).

**Figure 6.**
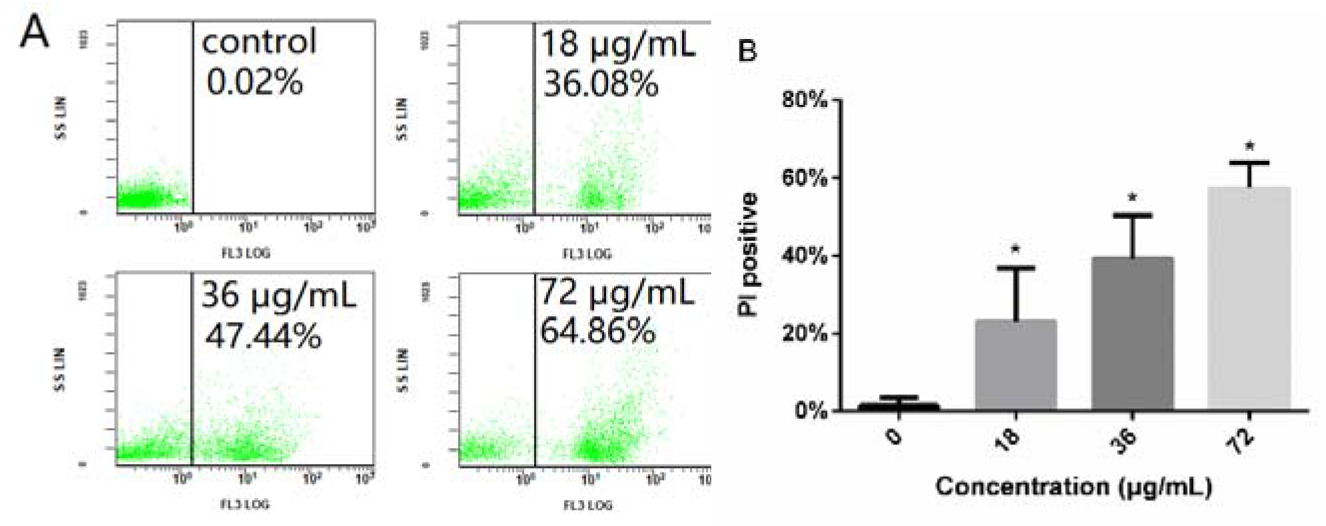
Ala induces permeability of plasma membrane of *C. albicans* cells. After exposure to 18, 36 and 72 μg/mL Ala for 4 h, fungal cells were stained with PI (final concentration: 10 μM) for 10 minutes before FCM analysis. A: representative FCM charts. B: statistical graph of PI staining. *, p<0.05, Ala-treated vs control.

The effects of Ala on ROS production in *C. albicans* cells were evaluated through FCM. As revealed by Figure 7, treatment with Ala significantly increased endogenous ROS production. Then, we assessed the role of ROS in the antifungal activity of Ala against biofilm formation. The addition of 5 mM NAC could save part of decreased viability of biofilms treated with 36 μg/mL of Ala, despite that 5 mM NAC alone could significantly reduce the viability of biofilms (Figure 8A). In addition, this kind of rescue by NAC could also be confirmed by microscope observations (Figure 8B), whereas the presence of NAC could restore the biofilm growth under the treatment of 36 μg/mL of Ala.

**Figure 7.**
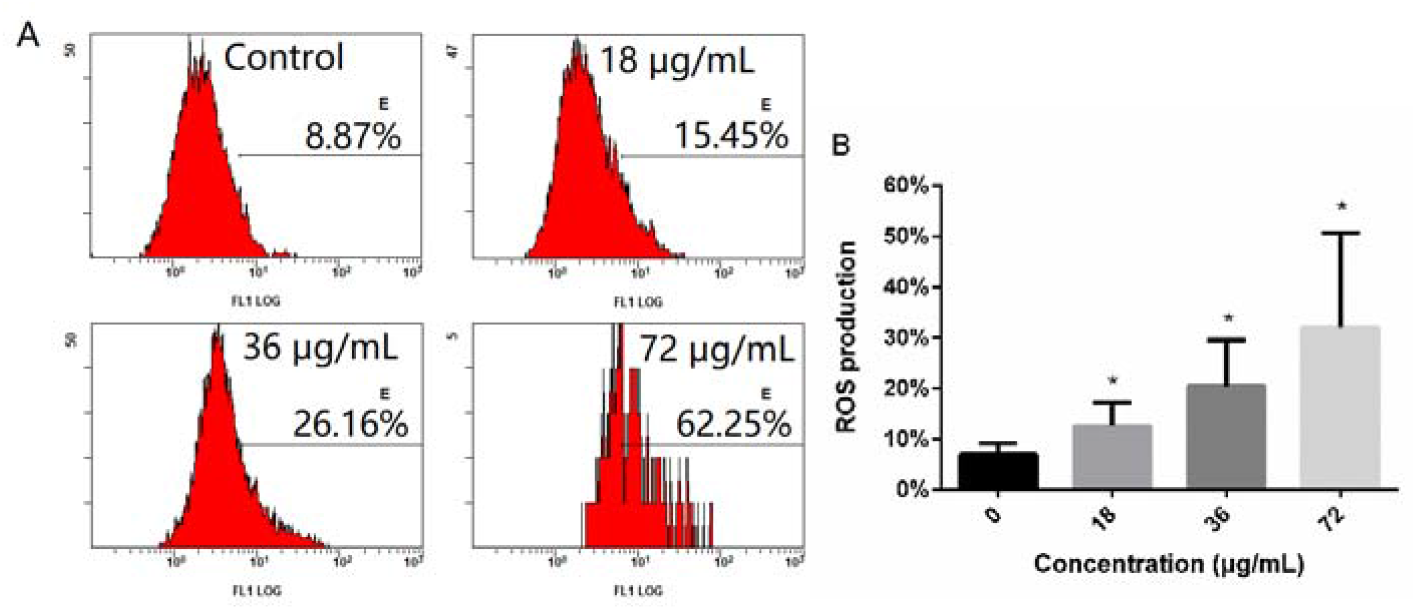
Ala increased endogenous ROS production in *C. albicans* cells. Fungal cells treated with different concentrations of Ala for 4 h were subjected to FCM to determine the intracellular ROS production. A: representative FCM charts. B: Statistical graph of ROS production. *, p<0.05 Ala-treated vs control.

**Figure 8.**
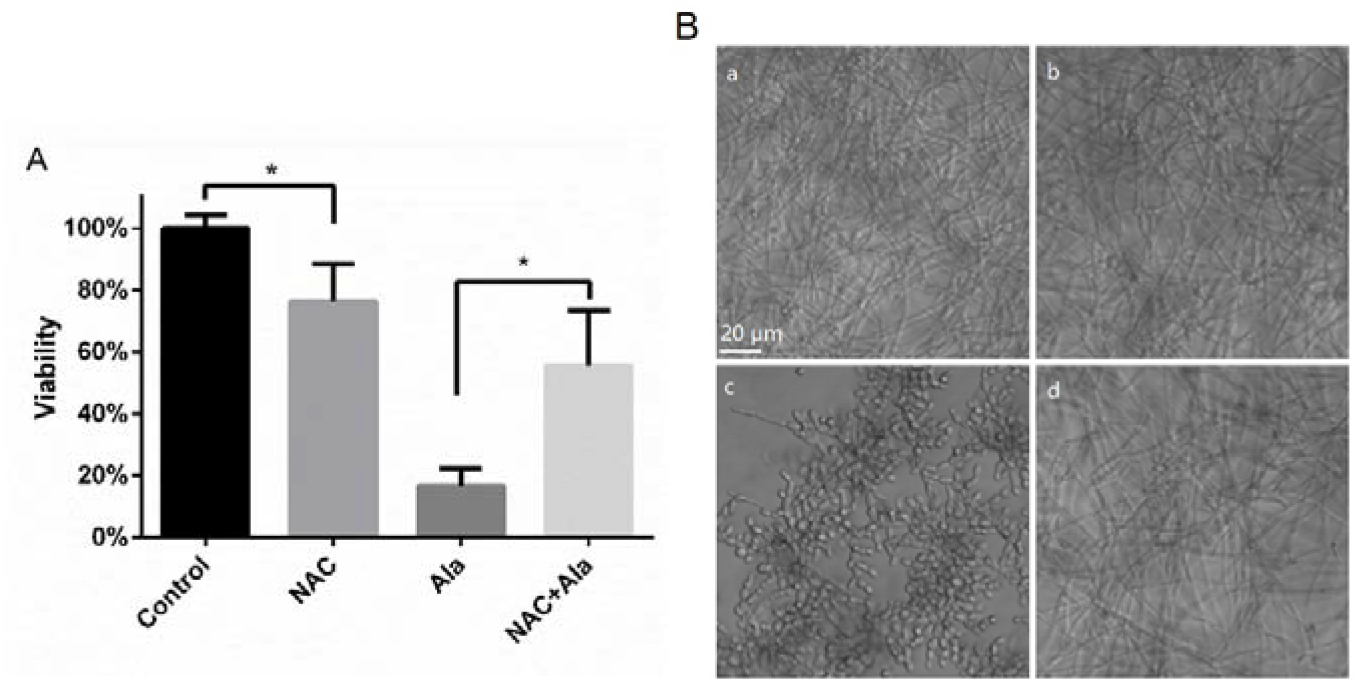
NAC could rescue the damages on biofilm viability imposed by Ala. Under biofilm-forming conditions, fungal cells in 1640 medium alone or with 5 mM NAC were allowed to grow 24 h in the absence or presence of 36 μg/mL Ala. XTT reduction assay were employed to determine the viability of each group (A) and the biofilm morphologies of each group were recorded by microscope (B). a, control; b, 5 mM NAC; c, 36 μg/mL Ala; d, 36 μg/mL Ala+5 mM NAC. *, p<0.05, determined by *t*-test.

To characterize the interactions between Ala and antifungal drugs, namely, AmB, FLZ and CAS, checkerboard assays based on microdilutions were performed. From data obtained, as shown in Table 2, the FIC of the combination of Ala and AmB was 0.75, suggesting an additive antifungal effect. In other words, Ala could potentiate the efficacy of AmB. Therefore, by adding Ala to the antifungal therapy, the dose of AmB could be decreased. This kind of additive and potentiating effect could also be seen in the combination of Ala and CAS. However, as for the combination between Ala and FLZ, the interaction was indifferent.

**Table 2.** Interaction of Ala with antifungal drugs

| Drug | MIC of drugs ($\mu\text{g/mL}$) | | MIC of Ala ($\mu\text{g/mL}$) | | FICI | Interaction |
| --- | --- | --- | --- | --- | --- | --- |
|  | Alone | Combined | Alone | Combined |  |  |
| AmB | 0.625 | 0.156 | 72 | 36 | 0.75 | Additive |
| CAS | 0.625 | 0.313 | 72 | 9 | 0.6125 | Additive |
| FLZ | 1.25 | 1.25 | 72 | 72 | 2 | Indifferent |

## Discussion

*C. albicans*, one of the most notorious human fungal pathogens, causes more than 250,000 death and millions of episodes of recurrent infections every year around the world, leaving a heavy burden on public health system (da Silva Dantas et al., 2016). The pressing situation we faced, including resistance and toxicity of currently available antifungal drugs, calls for new antifungal agents, especially agents effective against fungal biofilms which are highly resistant to antifungal agents and host defense (Wong et al., 2014; Lohse et al., 2018). In this study, we for the first time, to our knowledge, demonstrated the antifungal activity of Ala against *C. albicans*, as well as its biofilms and several other *Candida* species. Although this lactone has failed to show antifungal activities against *C. albicans* before, that may be due to the concentrations used (40 μg/mL), which was lower than the MIC of Ala in this study (Meng et al., 2001).

Biofilms formed on surfaces of medical devices and mucosal tissues represent a calcitrant reservoir for persistent infections, due to the complicated structures and resistance to antifungals (Mathe and Van Dijck, 2013). Adhesion constitutes the first prerequisite for *C. albicans* biofilm formation. Thus, many antifungal agents active against *C. albicans* biofilms showed suppression on adhesion, although biofilm-inhibitory agents without effects against adhesion did exist (Holtappels et al., 2018). In this study, Ala could inhibit the adhesion of *C. albicans* to polystyrene surfaces, upon which the biofilm formation of *C. albicans* could also be inhibited by Ala. More importantly, 18-72 μg/mL of Ala (1/4∼1 MIC) could exert disruptive effects on preformed biofilms, which could hardly be eradicated by antifungal drug fluconazole, as well as AmB at 1 MIC (Vila et al., 2013). This makes Ala a promising antibiofilm compound, although the MIC against *C. albicans* was a little high. Despite this, many factors present in vivo, such as liquid flow, host factors and components of immune response, may influence the biofilm formation and development (Nobile et al., 2012). The in vivo efficacy of Ala may also be influenced and warrant further investigation.

Yeast-to-hyphal transition is closely associated with the virulence of *C. albicans*, although this transition could be decoupled from pathogenicity (Noble et al., 2010). Multiple physical properties of *C. albicans* hyphae can be employed to promote active fungal invasion to epithelial tissues, especially those terminally differentiated (Richardson et al., 2019). Hyphae of *C. albicans* could produce candidalysin, a cytolytic peptide, to damage mucosal tissues, and hyphae can also facilitate the escape from macrophages and the formation of biofilms (Moyes et al., 2016; Noble et al., 2017). In the four kinds of hyphal-inducing media, Ala could inhibit the morphological transition, albeit that the degree of inhibition differed. The reason may be that the different signaling pathways were activated in those four kinds of media, among which the FBS-containing medium was the most potent inducer of hyphae (Sudbery, 2011). Meanwhile, even blocked in yeast state in the presence of 72 μg/mL Ala, *C. albicans* cells growing in FBS-containing SD medium outnumbered those growing in the other media. This is consistent with the fact that the presence of FBS could increase the MIC of CAS (Shields et al., 2011). Antifungal resistance of *C. albicans* biofilms may, at least partly, stem from the presence of EPS in biofilms, which could sequestrate the antifungal drugs (Taff et al., 2013). The complex physical and biological properties of EPS may warrant multitargeted or combination therapies (Karygianni et al., 2020). Ala could decrease the EPS of preformed biofilms, similar to antifungal agents active against biofilms published by others (Nithyanand et al., 2015). When used in combination with other antifungal drugs, Ala might facilitate the access of drugs to fungal cells within biofilms, thus posing a synergistic or additive effect on mature biofilms.

Extracellular phospholipase secreted by *C. albicans* contributes to the invasion and infection through degrading phospholipids in host cell membrane, while mutants of phospholipase showed less infectivity in experimental animal models of infection (Mayer et al., 2013; Singh et al., 2018). Many antifungal agents, such as fluconazole, dracorhodin perchlorate and 2, 4-di-tert-butylpheno, have shown inhibitory effects on phospholipase production (Padmavathi et al., 2015; Singh et al., 2018; Yang et al., 2018c). Similarly, in this study, the production of *C. albicans* extracellular phospholipase in the presence of Ala was less, in comparison to drug-free controls. Taken together, Ala could inhibit the growth and virulence factors of *C. albicans*.

Ala could significantly increase the portion of PI positive cells, suggesting that Ala permeabilized the plasma membrane. The increased permeability of *C. albicans* cell membrane can also be induced other antifungal agents, such as dioscin and hibicuslide C (Hwang et al., 2013; Yang et al., 2018b).

Among the mechanisms underlying antifungal agents and macrophages, ROS induction plays important roles (Erwig and Gow, 2016; Thangamani et al., 2017; Ries et al., 2019). Since Ala could induce ROS production in various kinds of tumor cells (Khan et al., 2012; Khan et al., 2013), so we speculated that ROS may be closely associated with the antifungal activity of Ala. As expected, Ala could significantly increase the production of ROS, consistent with our previous reports on Ala and other ROS-inducing antifungal agents (Khan et al., 2012; Khan et al., 2013; Thangamani et al., 2017; Yang et al., 2018a; Ries et al., 2019).

The famous antioxidant NAC showed inhibition on *C. albicans* growth and disruptive effects on preformed biofilms, although the effective concentrations were high, at the level of mg/mL (Venkatesh et al., 2009; Aslam and Darouiche, 2011). In our study, although 5 mM NAC (approximately 0.8 mg/mL) alone can inhibit the viability of *C. albicans* biofilms, which was consistent with its fungistatic property at high concentrations in previous researches by others (Aslam and Darouiche, 2011), supplement with 5 mM NAC could obviously rescue the decrease in viability caused by Ala treatment. This, along with the FCM results, confirmed that ROS did contribute to the antifungal activity of Ala. Combination therapy has been considered to extend the efficacy of current antifungal drugs and prevent/slow the emergence of drug resistance (Cui et al., 2015; Wright, 2016; Fisher et al., 2018). For example, the combinational use of fluconazole, flucytosine and AmB has been employed to treat cryptococcal meningitis in patients with HIV infections (Fisher et al., 2018)., the combination of Ala and AmB or CAS could produce an additive effect. This suggests that when used in combination, Ala might reduce the dose of AmB and CAS required to treat *C. albicans* infections.

## Conclusion

In conclusion, our current study reveals the antifungal effects of Ala on *C. albicans* for the first time. The virulence factors of this pathogenic fungus, including morphological transition, adhesion, biofilm formation, production of extracellular phospholipase, could be inhibited by Ala. Of note, the metabolic activity and EPS production of preformed biofilm also can be suppressed by this natural product. The antifungal effects of Ala against *C. albicans* may involve increased cell membrane permeability and ROS production.

## Availability of data and material

The all data could be obtained from the corresponding author.

## Authors’ contributions

Conception and design: XL, TM and LY; experiments: XL, LZ and LY, data analysis: XL, ZM and LY, review and comment: JX, ZM, TM and YS; All authors read and approved the manuscript.

The authors declare that all data were generated in-house and that no paper mill was used.

